# Molecular analysis of *Drosophila melanogaster* B chromosomes reveals their origin, composition, and structure

**DOI:** 10.1101/386995

**Authors:** Stacey L. Hanlon, Danny E. Miller, Salam Eche, R. Scott Hawley

## Abstract

The number of chromosomes carried by an individual species is one of its defining characteristics. Some species, however, can also carry supernumerary chromosomes referred to as B chromosomes. B chromosomes were recently identified in a laboratory stock of *Drosophila melanogaster*—an established model organism with a wealth of genetic and genomic resources—enabling us to subject them to extensive molecular analysis. We isolated the B chromosomes by pulsed-field gel electrophoresis and determined their composition through next-generation sequencing. Although these B chromosomes carry no known euchromatic sequence, they are rich in transposable elements and long arrays of short nucleotide repeats, the most abundant being the uncharacterized *AAGAT* satellite repeat. Fluorescent *in-situ* hybridization on metaphase chromosome spreads revealed this repeat is located on Chromosome *4*, strongly suggesting the origin of the B chromosomes is Chromosome *4*. Cytological and quantitative comparisons of signal intensity between Chromosome *4* and the B chromosomes supports the hypothesis that the structure of the B chromosome is an isochromosome. We also report the identification of a new B chromosome variant in a related laboratory stock. This B chromosome has a similar repeat signature as the original but is smaller and much less prevalent. We examined additional stocks with similar genotypes and did not find B chromosomes, but did find these stocks lacked the *AAGAT* satellite repeat. Our molecular characterization of *D. melanogaster* B chromosomes is the first step towards understanding how supernumerary chromosomes arise from essential chromosomes and what may be necessary for their stable inheritance.

## INTRODUCTION

The growth, development, and reproduction of an organism relies on genetic material that is organized into chromosomes. The chromosome complement carried by all members of a species is comprised of essential chromosomes referred to as the “A chromosomes.” A subset of individuals within a species may also possess extra chromosomes that are nonessential and not members of the standard A chromosome set. These supernumerary chromosomes are commonly referred to as “B chromosomes.” First described in 1907, B chromosomes have now been identified in hundreds of species across many different taxa, and it is estimated that B chromosomes may be present in 15% of all eukaryotic species (Wilson 1907; Jones and Rees 1982; Camacho 2005; D’Ambrosio *et al.* 2017).

The B chromosomes found in maize (*Zea mays*) are perhaps the best understood to date, and it is from their presence in corn that the name “Type-B chromosome” originated (Randolph 1928). The gravitation to study B chromosomes in corn was bolstered in part due to the advanced genetic and cytological tools available at the time, and B chromosomes have since served as useful tools to study and manipulate the genome (Beckett 1991; Birchler 1991; Yu *et al.* 2007; Masonbrink and Birchler 2012; Han *et al.* 2018). The recent rise of genomic analysis has allowed B chromosomes from several species to be examined on a molecular level, revealing more about their potential origin and genetic composition (Banaei-Moghaddam *et al.* 2015; Valente *et al.* 2017; Ruban *et al.* 2017). For instance, the sequencing of rye (*Secale cereal*) B chromosomes demonstrated they arose just over 1 million years ago and are derived from fragments of multiple A chromosomes (Martis *et al.* 2012). The B chromosomes in the cichlid fish *Astatotilapia latifasciata* were sequenced and found to carry euchromatic sequence— some of which is transcriptionally active—that is also present on several of the A chromosomes (Valente *et al.* 2014). More recently, sequencing the B chromosome of the grasshopper *Eumigus monticola* indicated that while it likely arose from one of the A chromosomes, it has since undergone amplification (Ruiz-Ruano *et al.* 2017), and deep-sequencing in another grasshopper, *Eyprepocnemis plorans*, revealed the presence of 10 protein-coding genes carried on the B chromosome (Navarro-Domiínguez *et al.* 2017). These studies demonstrate that the diversity and complexity of supernumerary B chromosomes has been underestimated, and a better understanding of their origin and composition is necessary.

Further work on the molecular characterization of B chromosomes has been impeded by several obstacles. First, the majority of identified B chromosomes have only been studied cytologically in samples collected from wild populations. Many of these species are not conducive to husbandry in the lab, precluding development of the molecular, genetic, and genomic tools necessary to allow in-depth study of B chromosomes. Second, the inability to control breeding and environment has the potential to introduce uncontrolled genetic variation and unknown pressure on B chromosome evolution, leading to fluctuations in B chromosome variants, frequency, and transmission rates (Zurita *et al.* 1998; Araújo *et al.* 2002; Bakkali and Camacho 2004; Manrique-Poyato *et al.* 2013; Lanzas *et al.* 2018). Third, it is challenging to precisely determine the age of a B chromosome in these wild systems, making it difficult to discern how they formed and what events led to their current composition. Thus, as pointed out by Jones (1995), B chromosome observations are only able to reflect the system in the present and therefore may not reflect how the system was in the past.

Recently, B chromosomes were identified in a laboratory stock of *Drosophila melanogaster* (Bauerly *et al.* 2014). Although B chromosomes have previously been observed in a handful of wild populations within the *Drosophila* genus [*albomicans* (Clyde 1980); *malerkotliana* (Tonomura and Tobari 1983); *kikkawai* (Sundaran and Gupta 1994); *subsilvestris* (Gutknecht *et al.* 1995); *lini* and *pseudoananassae* (Deng *et al.* 2007)], Bauerly *et al.* (2014) was the first report of B chromosomes in *melanogaster.* The laboratory stock in which these were identified carries a null allele of the female germline-specific gene *matrimony* (mtrm). This allele, referred to as *mtrm^126^*, was generated through the imprecise excision of a P-element nearly 10 years earlier (Xiang *et al.* 2007). Presently, the number of B chromosomes in this *mtrm^126^* stock averages 10–12 copies in addition to the A chromosome complement (Figure 1A and B). Bauerly *et al.* (2014) demonstrated these B chromosomes have centromeres since they incorporate the *Drosophila* centromeric histone variant CID and were able to interact with the meiotic spindle. They also analyzed female meiotic prometaphase chromosomes using fluorescent *in situ* hybridization (FISH) and showed that the B chromosomes carry the *AATAT*satellite sequence, which is found on the *X* and *Y* chromosomes but is primarily on Chromosome *4*. Complementation tests and qPCR indicate the B chromosomes do not carry euchromatic material from Chromosome *4*, but they do carry large amounts of heterochromatin due to the abundance of histone H3 trimethylation on K9 (H3K9m3) and their pronounced effect on position effect variegation (Bauerly *et al.* 2014).

**Figure 1.**
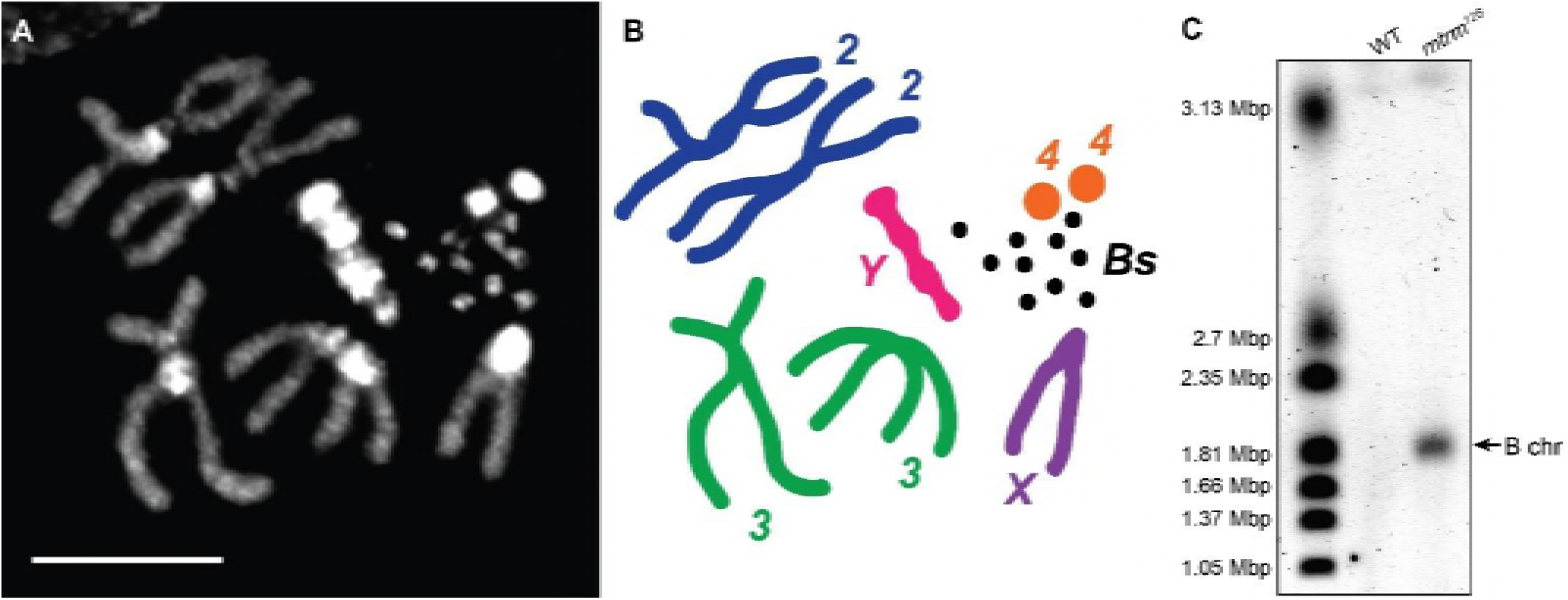
Cytological and molecular evaluation of the *D. melanogaster* B chromosomes. (A) Metaphase chromosome spread of a male with 10 B chromosomes, collected from the *mtrm^126^* stock that carries an average of 10–12 B chromosomes. Scale bar = 5 μm. (B) Illustrated representation of the karyotype in (A). (C) Results of PFGE after gel was stained with GelRed (image is inverted). The B chromosomes form a single, distinct band around 1.81 Mbp that is absent in the wild-type (WT) stock. Size standard: *H. wingeii* chromosomal DNA (Bio-Rad).

Several questions loom large regarding the origin and composition of the *D. melanogaster* B chromosomes. It is presently unclear if the B chromosomes are homogeneous and/or if new B chromosomes are continually forming from the A chromosomes. Current cytological methods of B chromosome examination cannot resolve subtle differences in size that may exist between variants that arose at different times. For instance, if the B chromosomes are all the same size, it is likely there was a single creation event followed by extensive proliferation, whereas an array of B chromosome sizes is more consistent with either multiple creation events or overall B chromosome instability. Additionally, the genetic composition of the B chromosomes is poorly understood: it is not clear what, if any, euchromatin the B chromosomes possess, and the known content of its heterochromatin is limited to the *AATAT* satellite repeat.

Here, we report the purification and sequencing of the *D. melanogaster* B chromosome described in Bauerly *et al.* (2014). Deep sequencing of this B chromosome reveals no known protein-coding genes but does show that these B chromosomes contain several highly repetitive elements that map to unplaced scaffolds from the current *D. melanogaster* genome assembly. One of these elements, the previously uncharacterized *AAGAT* satellite repeat, was used as a FISH target on metaphase chromosome spreads. We find this repeat to be specific to Chromosome *4* as well as enriched twofold on the B chromosomes, suggesting that the B chromosome is likely an isochromosome that arose from Chromosome *4*. Surprisingly, FISH analysis of a related stock that carries a different null allele of *mtrm* revealed the presence of a second, smaller B chromosome. Although present at a much lower copy number, this newly identified B chromosome variant has the same suite of satellite repeats carried by the original B chromosome, including the *AAGAT* satellite. Our characterization of these two B chromosomes on a molecular level provides the start of a better understanding of how new chromosomes may arise and become essential fixtures of the genome.

## MATERIALS AND METHODS

### Stocks used in this experiment

All stocks used in this report can be found in Table S1. Stocks were maintained on standard cornmeal media supplemented with an additional sprinkle of active dry yeast and kept at 24° and 70% humidity in constant light conditions.

### Sample collection for pulsed-field gel electrophoresis (PFGE)

To obtain samples for PFGE, 20 L3 larvae were placed in 3 mL Ringer's solution in a 35-mm glass petri dish. The brain and any attached imaginal discs were dissected from each larva one at a time, and care was taken to ensure the desired tissue was treated as gently as possible to avoid trauma. The tissue was transferred to a 1.5-mL microfuge tube with 500 μL Ringer’s solution kept on ice while the remaining dissections took place. Dissections of 20 larvae took no longer than one hr (typically 30–40 min). Tissue was briefly spun down to the bottom of the tube, the Ringer’s solution was removed, and 30 μL of insert buffer (0.1 M sodium chloride, 30 mM TrisCl pH 8, 50 mM EDTA pH 8, 0.5% Triton X-100, 7.7 mM 2-mercaptoethanol) was added. The tissue was homogenized by hand-grinding with a plastic pestle for 10 circular strokes. The tube was briefly spun to get the material down, then the homogenization was repeated twice more to ensure complete homogenization. The tube was briefly spun down for a final time and the top-most 25 μL was moved to a new tube in order to minimize the amount of cellular debris included in the plug. This new tube was placed at 42° for 2 min to warm up. Using a cut-off pipette tip (1 mm inner diameter), 75 μL of molten 1% InCert agarose (in 0.125 M EDTA) held at 50° was added to the warmed homogenate and mixed by pipetting gently three times, followed by distribution into a mold for a single plug. The mold was placed at 4° to set and remained there while additional rounds of dissections took place. After dissections were finished, plugs were transferred to 50-mL conical vials (no more than 4 plugs/vial) with 2.5 mL/plug NDS (0.5 M EDTA, 10 mM TrisCl pH 8, 1% sarkosyl, pH 9.5 and sterile-filtered) and 0.1 mg/mL Proteinase K. The tube with the plugs was incubated in a 50° water bath for 12.5 hr and gently swirled once or twice during the incubation. After the Proteinase K treatment, the tubes were placed at 4° until processing, which was usually later the same day (1–2 hr after the end of the incubation).

To process the plugs, they were moved from the NDS+Proteinase K solution into new conical tubes with 3 mL/plug 1x TE (pH 7.5) with 0.1 μM PMSF. Tubes were gently agitated on an orbital shaker for 1 hr at room temperature, then the 1x TE + PMSF was replaced with fresh 1x TE + PMSF and the tubes were shaken for another hour at room temperature. This was followed by four washes with 1x TE for 20 min each at room temperature, after which the plugs were moved into 1x TAE.

### PFGE setup and parameters

To set up the pulsed-field gel rig, 1x TAE was circulated and chilled to 14°. The gel was made with 0.8% SeaKem Gold agarose (in 120 mL 1x TAE) that was held at 50° until poured. Sample plugs as well as two plug standards with chromosomes from *H. wingei* (BioRad) were arranged on the front of a gel comb so they would be embedded in the gel. Approximately 100 mL of the gel was poured and allowed to set for 15 min at 4°. Once the gel was set, the comb was removed, and the empty wells were filled with the remaining unpoured gel and left to set for 5 min. The gel was then placed in the apparatus, followed by the start of the run with the following parameters: 500 s pulse time, 106° included angle, 3 V/cm with linear ramp, 45 hr total run time.

### B chromosome DNA Isolation

Initially, after the PFGE run (see above), the entire gel was stained in 3x GelRed and imaged to determine the size of the B chromosomes. Once their position was reliably determined, additional PFGE runs were conducted and the B chromosome DNA was extracted as follows. The ladder lanes were cut off and stained for 30 min in 0.5x SYBR Gold in 1x TAE in order to provide a guide for where the B chromosomes were located in the gel. The portion of the gel in the 1.81-Mbp range (where the B chromosomes run) was excised and the remainder of the gel stained to verify the correct part of the gel containing B chromosomes was removed.

To extract the B chromosome DNA, the excised gel fragment was placed in dialysis tubing (MWCO = 12K–14K) that was clipped closed at both ends and a second round of electrophoresis was conducted in a normal gel rig. This enables the B chromosome DNA to migrate out of the gel and into the buffer contained within dialysis tubing. Once complete, the buffer containing the B chromosome DNA was pipetted from the tubing and transferred to a 1.5-mL microfuge tube and dried down to concentrate the sample.

### DNA library creation and sequencing

The B chromosome DNA sample was sonicated with a Covaris 220 to ~ 600-bp fragments and proceeded directly to library construction using the KAPA High Throughput Library Prep Kit (KK8234), and the Bioo Scientific NEXTFlex DNA Barcodes (NOVA-514104) for barcoding. Quality control was completed using the Agilent 2100 Bioanalyzer and the Invitrogen Qubit 2.0 Fluorometer. The sample was sequenced on the Mi-Seq via a Mi-Seq 150-bp paired-end run. This run used MiSeq Reporter Control Software (v2.6.2.3) and RTA version MCS 2.6.1.1.

### Genome alignment, assembly, and analysis

Reads were aligned to dm6 using bwa (Li and Durbin 2009). Unmapped reads were identified by ‘samtools view -f4’ and unmapped read pairs were isolated and trimmed with sickle (https://github.com/najoshi/sickle) and scythe (https://github.com/ucdavis-bioinformatics/scythe), removing bases with quality scores < 30 and excluding any read < 40 bp after trimming. SOAPdeNovo2 was used to assemble unmapped reads using a kmer size of 41 and average insert size of 300 (Luo *et al.* 2012). Assembly generated 347,712 contigs, 75 of which were larger than 500 bp with a maximum contig size of 1,793 bp. To determine whether the 75 contigs larger than 500 bp contained protein-coding sequence, BLASTx (Altschul *et al.* 1990) was used to search RefSeq (update date 2018-04-02, containing 103,725,652 sequences) (O’Leary *et al.* 2016).

### Chromosome squashes

#### For DNA staining and FISH

For chromosome spreads from larval brain tissue, a protocol adapted from Larracuente and Ferree (2015) was used. Before beginning, 2 mL of fixative solution (45% acetic acid, 2.5% paraformaldehyde) was made fresh. A single larva was removed and placed in a fresh 50-μL drop of 0.7% sodium chloride. The brain and associated imaginal discs were extracted in the same fashion as for the PFGE (see above). The brain was then moved to a fresh 50-μL drop of 0.5% sodium citrate for hypotonic treatment for 5 min, followed by a transfer to the fixative solution for 4 min. After fixation, the brain was transferred to a 3-μL drop of 45% acetic acid on an 18-mm x 18-mm No. 1.5 siliconized coverslip. A microscope slide was inverted onto the coverslip with the sample and pressed gently to spread the liquid to the edges of the coverslip. The slide+coverslip was squashed for 2 min using a hand clamp (Milwaukee Tools, 48–22–3002), then immediately placed into liquid nitrogen for at least 5 min. Immediately after removal, the coverslip was popped off the slide using a razor blade. The slide was dehydrated by placing it in 70% ethanol for at least 10 min at -20°, then transferred to 100% ethanol at -20° for at least 10 min. Slides were removed and allowed to completely air-dry, then stored in a corked slide box kept at room temperature.

For chromosome spreads from ovary tissue, the protocol used for larval brains described above was used with the following changes. Ovaries were collected from mated adult females by anesthetizing them with carbon dioxide until incapacitated (~5–10 seconds). A single female was then moved to a fresh 50-μL drop of 0.7% sodium chloride and whole ovaries were removed. The tips were separated from the later stages such that anything larger than stage 8 was discarded. The ovary tips were then hypotonically treated and fixed as above for larval brains. Before squashing, the tips were gently teased apart on the siliconized coverslip in 3 μL 45% acetic acid in order to spread out the tissue. Squashing, freezing, dehydration, drying, and storage were all carried out as described above for larval brains.

#### For immunofluorescence (IF)

To preserve protein-DNA interactions in metaphase chromosome spreads from larval brain tissue, a protocol similar to the acid-free squash technique in Johansen *et al.* (2009) was used. Before beginning, 2 mL of Brower’s fixative solution (0.15 M PIPES, 3 mM magnesium sulfate, 1.5 mM EGTA, 1.5% Nonidet P-40 substitute, 2% paraformaldehyde) was made fresh. Larval brains were removed in 0.7% sodium chloride and incubated in 2 mL 50 μM colchicine in 0.7% sodium chloride at 25° for 1 hr. (The colchicine arrests cells in metaphase, resulting in super-condensed chromosomes that are able to hold their shape better after Brower’s fixative.) After the incubation, the brain was transferred to a fresh 50-μL drop of 0.5% sodium citrate for hypotonic treatment for 5 min, followed by a transfer to the fixative solution for 2 min. The brain was then moved to a 50-μL drop of 50% glycerol for 5 min, then to 5-μL drop of 50% glycerol on an 18-mm x 18-mm No. 1.5 siliconized coverslip. The brain tissue was gently dissociated using a bent needle probe, then a microscope slide was inverted onto the coverslip and direct pressure was applied to spread the tissue. The slide+coverslip was squashed, frozen, and dehydrated as described above. Slides were allowed to air-dry for 10 min, followed by immediate processing for IF.

### Sample slide processing

#### For DNA staining and FISH

For simple DNA visualization, 5 μL of Vectashield with DAPI mounting media was applied to a clean 22-mm x 22-mm No. 1.5 glass coverslip that was placed on the sample slide,then sealed to the slide with clear nail polish.

For FISH on the chromosome spreads, a protocol adapted from Larracuente and Ferree (2015) was used. For each slide, 21 μL of the FISH solution (50% formamide, 10% dextran sulfate, 2x SSC, 100 ng fluorescently-labeled probe (see Table S2) was applied directly to the dried slide and a clean 22-mm x 22-mm No. 1.5 glass coverslip was placed on top. Gently, any large bubbles that in contact with the tissue were nudged to the edges. The slide was heated to 95° on a heat block for 5 min in darkness, then transferred to a container lined with damp paper towels (to maintain humidity and prevent the samples from drying out) and placed at 30° overnight (16–24 hr). After the incubation, slides were washed three times in 0.1x SSC for at least 15 min each. Slides were blown dry, then mounted by applying 5 μL Vectashield with DAPI to a clean 22-mm x 22-mm No. 1.5 glass coverslip that was placed on the sample slide, then sealed to the slide with clear nail polish.

#### For IF

Dried slides were placed in PBSTX (1x PBS with 0.1% Triton X-100) and washed three times for 5 min each. Slides were then blocked with 5% dry, nonfat milk in PBSTX for 20 min at room temperature. Excess block was wiped away from the perimeter of the sample area and a primary antibody solution (antibody in 30 μL block solution; 1:500 rat anti-CID from a test bleed generated for the Hawley lab, 1:100 rabbit anti-HOAP serum, a gift from Y. Rong) was applied, and a 22-mm x 22-mm No. 1.5 glass coverslip was placed on top. The slides were placed in a container lined with damp paper towels and placed at 4° overnight (16–24 hr). After the incubation, the slides were washed with PBSTX three times for 5 min each. Excess PBSTX was wiped away from the perimeter of the sample area and a secondary antibody solution (antibody in 30 μL block solution; 1:500 anti-rat from Invitrogen, 1:500 anti-rabbit from Invitrogen) was applied, and a 22-mm x 22-mm No. 1.5 glass coverslip was placed on top. The slides were placed in a container lined with damp paper towels and placed in a dark drawer at room temperature for 45 min. After the incubation, the slides were washed with PBSTX three times for 5 min each. Excess PBSTX was wiped away from the perimeter of the sample area and 3 μL Vectashield with DAPI was applied to a clean 22-mm × 22-mm No. 1.5 glass coverslip that was placed on the sample slide, then sealed to the slide with clear nail polish.

### Microscopy and image processing

Images were acquired with a DeltaVision microscopy system (GE Healthcare, Piscataway, NY) consisting of a 1×70 inverted microscope with a high-resolution CCD camera. All imaging used a 100× objective and 1.6× auxillary magnification. Stacks of 15 *z* images with a thickness of 0.2 μm were taken for each metaphase using 100% transmission and standard DAPI, FITC, and TRITC filters. Images were deconvolved using SoftWoRx v. 6.1.3 or later (Applied Precision/GE Healthcare) following Applied Precision protocols. Stacks of deconvolved images were combined in a z projection showing maximum intensity, cropped to the region of interest, recolored, and adjusted for brightness and contrast in FIJI/ImageJ.

For direct comparisons of *AAGAT* FISH signal, samples were collected, prepped, and imaged concurrently. Settings for excitation/emission/exposure were identical for all four samples. Stacks of deconvolved images were combined in a *z* projection showing maximum intensity, cropped to the region of interest, and recolored in FIJI/ImageJ. Only the DAPI channel was adjusted for brightness and contrast; no adjustments were made to the FITC and TRITC channels, thus the display range of each image in those channels represents the full, original pixel value range.

### Intensity measurements of DAPI and FISH probes

After imaging (as described above), deconvolved image stacks were opened in FIJI/ImageJ and made into a z projection by summing the intensity of the slices in the stack (instead of maximum intensity projections) in order to preserve the total intensity. A circular region of a defined size was drawn around chromosomes (either B1, B2, or Chromosome *4*) that were well separated from other chromosomes. An equal number of regions were placed near the circled chromosomes in order to obtain background intensity values.

Integrated density measurements from both the DAPI and FISH channels within each region were taken, and measurement values from like chromosomes were averaged for each metaphase individually; for example, each of the measurements for the B1 chromosomes in a single metaphase were averaged, as well as the values for the background measurements within the same single metaphase. When there was only one of a chromosome, the measured value was used. To adjust for the background, the average of the background measurements was subtracted from the average (or single value) of the chromosomes. These adjusted values were then divided by one another in order to get the appropriate statistic.

### Data availability and Supplemental Material

Illumina sequencing reads generated in this report are available at the National Center for Biotechnology Information (https://www.ncbi.nlm.nih.gov/) under project PRJNA484270. Scripts used in the analysis of this project can be found at https://github.com/danrdanny/B-chromosome/. Original data underlying this manuscript can be accessed from the Stowers Original Data Repository at http://www.stowers.org/research/publications/LIBPB-1334. Supplemental files available at FigShare. Supplemental File S1 includes all supplemental figures (Figures S1–4) and tables (Tables S1–4). Supplemental File S2 are blast results. All stocks and reagents available upon request.

## RESULTS

### Molecular isolation of *D. melanogaster* B chromosomes

The B chromosomes reported in Bauerly *et al.* (2014) were first identified in a stock carrying a null allele of matrimony (*mtrm^126^*) generated by the Hawley laboratory around 2004. Matrimony is a Polo-kinase inhibitor that is specific to female meiosis (Xiang *et al.* 2007). The level of Mtrm, relative to Polo kinase, is important during oogenesis: half the wild-type amount of Mtrm results in the missegregation of noncrossover chromosomes during meiosis I, and the complete absence of Mtrm leads to meiotic catastrophe and sterility (Bonner *et al.* 2013). Individual flies within this *mtrm^126^* stock possess an average of 10–12 B chromosomes as assessed by examining metaphase chromosomes from brains of third-instar larvae (Figures 1A and B).

Due to the increased potential for an aberrant female meiosis in this stock, we wondered if the B chromosomes were the product of a chromosomal fragmentation event (or events) that may have occurred following a *mtrm*-induced premature entry into meiotic anaphase I (Bonner *et al.* 2013). If the present B chromosomes are copies of an original single, stable chromosome fragment, then we expect all B chromosomes to be the same size, whereas if each B chromosome was a product of a different fragmentation event, each one would likely be a different size. Because the current cytological method of B chromosome analysis is not sensitive enough to detect small size differences between B chromosomes, we looked for a method that would allow a more precise size measurement of the B chromosomes. Pulsed-field gel electrophoresis (PFGE), a technique routinely used to resolve large DNA fragments in the megabase range, enabled us to measure the size(s) of intact B chromosomes. Brains and imaginal discs from male and female third-instar larvae in the *mtrm^126^* stock were disrupted and embedded in agarose, with all proceeding treatments occurring in-gel to prevent shearing of whole chromosomes (see Materials and Methods). Our PFGE produced excellent resolution within the approximate size range of the B chromosome, which appeared as a single, distinct band around 1.81 Mbp (Figure 1C). We did not see a smear on the gel that would represent B chromosomes of various sizes, nor did we observe additional bands that may represent variants of B chromosomes that may have arisen independently. The presence of an individual, distinct band leads us to conclude that the B chromosomes in this stock are of very similar size, and accordingly, we favor a model where an individual B chromosome proliferated within the *mtrm^126^* stock to reach the present high copy number.

### Sequence analysis of isolated B chromosomes confirms their heterochromatic composition

Isolation of the *D. melanogaster* B chromosomes by PFGE gave us an opportunity to study them in the absence of the standard chromosome complement. After PFGE, the portion of the gel encompassing the B chromosomes was removed and the DNA extracted, purified, and sequenced via paired-end Illumina sequencing (see Materials and Methods). We generated 42,964,020 150-bp paired-end reads (21,482,010 read pairs) and aligned them to release 6 (dm6) of the *D. melanogaster* genome. A total of 30% (13,031,938) of all reads aligned in pairs to unique locations in the reference genome, while 33% (14,212,886) of reads mapped to either non-unique locations or were pairs that did not align properly. The remainder of the reads (37%, 15,719,196) did not map to known genomic locations.

Among the 13,031,938 uniquely mapping reads, 84% aligned to the *X*, *2^nd^, 3^rd^*, or *Y* chromosomes with an average depth of coverage of 12x (Table S3). We anticipated this low, evenly distributed depth of coverage across the genome since our PFGE purification would likely include a random assortment of genomic fragments that were in the size range of the B chromosome. We then masked repetitive sequence using RepeatMasker and plotted depth of coverage to identify regions that exceeded this background (Smit *et al.* 2015). We found no focal increases in coverage along the euchromatic regions of the genome, confirming that the B chromosomes do not carry any euchromatic genomic sequence (Figure S1). As for Chromosome *4*, 5% of the uniquely mapping reads aligned with an average depth of coverage of 76x. Masking repetitive sequence reduced this to approximately 10x, similar to the genome-wide background. This reduction in coverage demonstrates that many reads that uniquely mapped to Chromosome *4* are in repetitive or low-complexity areas. This observed lack of euchromatic sequence in our isolated B chromosome sample confirms previous work that suggested the B chromosomes are highly heterochromatic (Bauerly *et al.* 2014).

The remaining 11% of uniquely mapped reads aligned to small reference scaffolds whose exact position in the genome is unknown. These unplaced scaffolds are generally short, highly repetitive, or low complexity fragments, suggesting they are heterochromatic sequences. Of 1,870 scaffolds present in the dm6 assembly, we found 133 that contained at least one region where 100 or more reads uniquely aligned, suggesting these scaffolds contain sequence present on the B chromosomes. In total, 114,673 bp of total sequence across these 133 contigs were covered by at least 100 reads (Table S4). As an example, the 51-kb scaffold chrUn_DS483562v1 had at least 100 reads align uniquely over more than 80% of its sequence, suggesting that it contains a large, but not necessarily contiguous, amount of DNA present on the B chromosomes.

We also wondered if the B chromosomes carried any novel sequence—euchromatic or heterochromatic—not present in the current release of the Drosophila genome. To determine if any of the 15,719,196 unmapped reads were from unique, non-repetitive sequences not represented within the *D. melanogaster* genome assembly, we performed a *de novo* assembly using SOAPdenovo2 (Luo *et al.* 2012) using a kmer size of 41 (see Materials and Methods). This resulted in the assembly of 75 contigs greater than or equal to 500 bp with an average assembled length of 744 bp (max 1,793 bp). We then searched RefSeq using BLASTx to determine if these contigs contained any complete genic sequences or other known sequence and found that 39 of 131 contigs contained at least one significant hit in the RefSeq database. Of all the proteins identified using our method, none were complete copies and the majority were viral, suggesting that several of these contigs contained fragments of transposable elements (File S2). Taken together, this sequence analysis suggests that the B chromosomes we sequenced are derived from the *D. melanogaster* genome and do not contain any previously known or novel euchromatic sequence.

### FISH analysis indicates Chromosome *4* is the source of the B chromosomes

Previous analysis of female meiotic prometaphase chromosomes using FISH revealed the B chromosomes carry the simple nucleotide repeat *AATAT*. Though the B chromosomes are able to move independently during prometaphase, the remainder of the chromosomes form a mass that makes it difficult to discern where a repetitive sequence resides on an individual chromosome. Additionally, since the *Y* chromosome is absent in female prometaphases, we cannot perform extensive FISH analyses to rule it out as a potential source of the B chromosomes. To circumvent these obstacles, we performed FISH using the same *AATAT* probe on metaphase chromosomes from male third-instar larval brains. We were able to verify the previously described presence of this repeat on the B chromosomes, as well on Chromosome *4*, the heterochromatic tip of the *X*, and several locations on the *Y* chromosome (Lohe *et al.* 1993; Bauerly *et al.* 2014) (Figure 2A).

**Figure 2.**
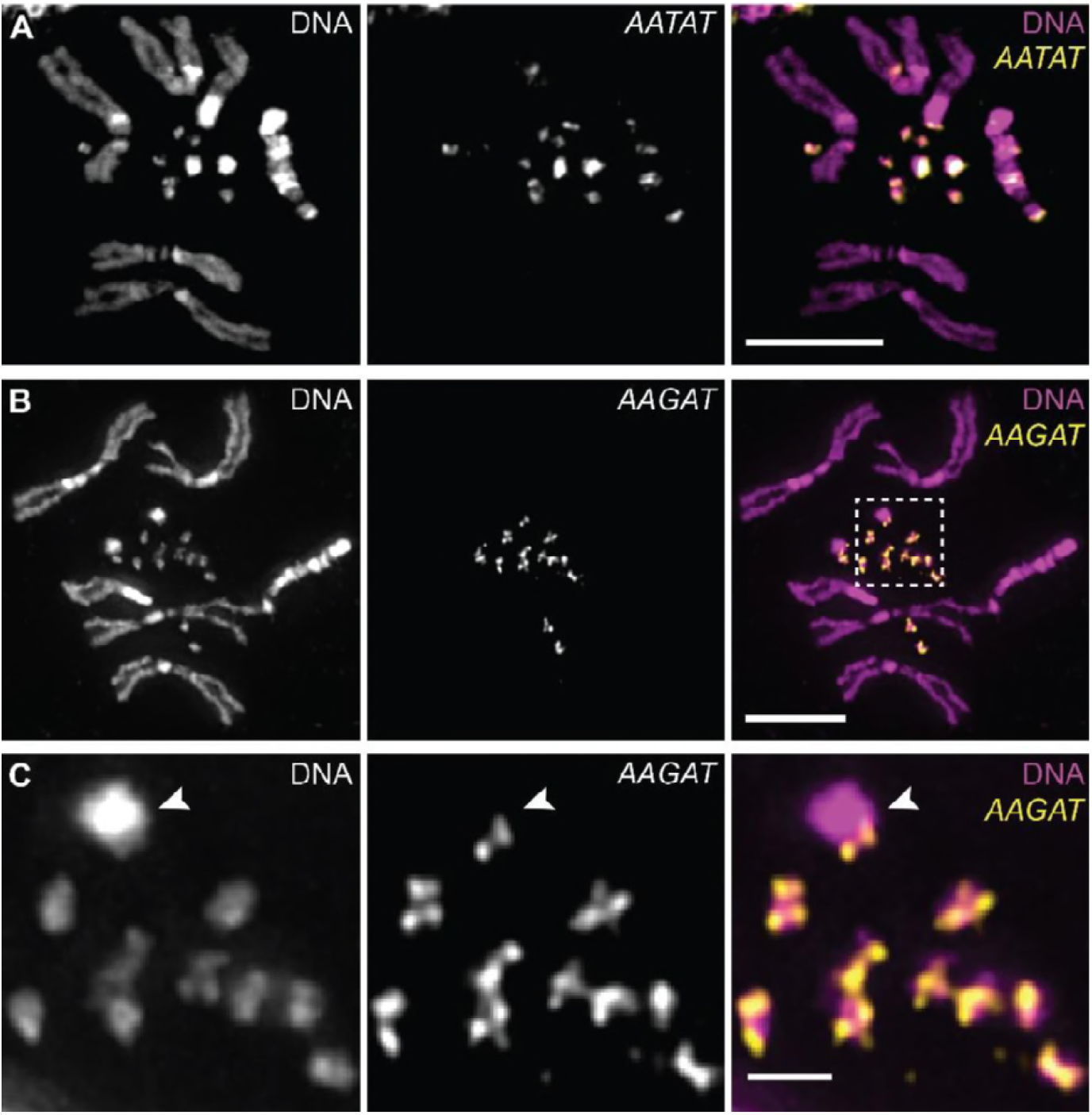
FISH on metaphase chromosome spreads using probes that recognize simple satellite repeat sequences. (A) A probe recognizing the *AATAT* repeat hybridizes to the B chromosomes, Chromosome *4*, the tip of the *X* chromosome, and various locations on the *Y* chromosome. Scale bar = 5 μm. (B) A probe recognizing the *AAGAT* repeat hybridizes only to the B chromosomes and Chromosome *4*, indicating the B chromosomes arose from Chromosome *4*. Scale bar = 5 μm. (C) Magnification of Chromosome *4* (arrowhead) and the B chromosomes from the box in (B). Scale bar = 1 μm.

Since the B chromosome may have arisen from any of these chromosomes, we wondered if our B chromosome sequence contained short, tandemly repeating kmers that could be used as FISH targets. We created an analysis pipeline modeled after one used previously (Wei *et al.* 2014) to generate a list of kmers present in the B chromosome raw sequencing data (Table 1). Among the kmers with over a million counts, we recovered the *AATAT* repetitive sequence, confirming our previous FISH results. Additionally, we found the *AAGAG* repetitive sequence was enriched on the B chromosomes as well as Chromosome *4* (Figure S2A), however this repeat can also be found on each of the other chromosomes as it is one of the most abundant 5mer repeats in the genome (Lohe *et al.* 1993; Wei *et al.* 2014). Other repetitive sequences that are abundant in genome-wide studies but were not prevalent in our kmer analysis do not appear to be present on the B chromosomes (Figure S2B-F). The most prevalent repetitive sequence identified in our kmer analysis was the 5mer *AAGAT*, which has appeared recently in two genomic sequencing datasets but has yet to be fully characterized (Wei *et al.* 2014; Talbert *et al.* 2018). Using a FISH probe for the *AAGAT* repeat, we found it to be highly enriched on the B chromosomes as well as at the tip of Chromosome *4* (Figures 2B and 2C). As Chromosome *4* is the only chromosome that shares all three of the most abundant repeats found on the B chromosome, it is likely to be the source of the B chromosomes.

**Table 1.** List of the most abundant kmers from our sequence analysis of the B chromosome.

| Kmer | Count | % of maximum | kmer category |
| --- | --- | --- | --- |
| TAAGA | 40,607,724 | 100% | AAGAT |
| GATAA | 31,022,996 | 76% |  |
| ATAAG | 27,013,024 | 67% |  |
| AAGAT | 16,852,053 | 41% |  |
| AGATA | 16,545,657 | 41% |  |
| TATAA | 10,971,510 | 27% | AATAT |
| TAATA | 10,414,421 | 26% |  |
| AATAT | 9,354,811 | 23% |  |
| ATATA | 7,010,495 | 17% |  |
| ATAAT | 4,812,487 | 12% |  |
| A | 4,235,133 | 10% | A |
| AGAAG | 3,402,400 | 8% | AAGAG |
| AAGAG | 2,840,917 | 7% |  |
| GAAGA | 2,743,196 | 7% |  |
| AGAGA | 2,652,461 | 7% |  |
| GAGAA | 2,172,940 | 5% |  |

### B chromosomes have functional telomeres that recruit the capping protein HOAP

Due to its smaller size and its similarity in composition, the initial B chromosome may have been a fragment that broke away from Chromosome *4*. A double-strand break on Chromosome *4* would produce two fragments, each with a single telomere and a single broken end. Only one of these fragments, however, would possess the centromere and be able to be segregated during cell division. For the centric fragment to be adequately maintained, its broken end would need to be mitigated, such as through the formation of a ring chromosome that would negate a need for telomeres altogether (Titen and Golic 2008). Alternatively, the broken end may become a functional neo-telomere that can recruit telomeric capping proteins, which has previously been observed in *Drosophila* (Gao *et al.* 2010; Kurzhals *et al.* 2017).

To distinguish between these possibilities, we wanted to determine if the B chromosomes have functional telomeres by looking for the presence of the telomere-specific capping protein HOAP (encoded by cav) on chromosomes in metaphase spreads (Cenci *et al.* 2003). Though the protocol used for FISH analyses produces excellent chromosome morphology, acetic acid in the fixative solution is known to remove proteins associated with DNA, such as histones (Dick and Johns 1968; Pimpinelli *et al.* 2011). To avoid this, we switched to a protocol similar to the acid-free squash technique in Johansen *et al.* (2009). Using antibodies that recognize the *Drosophila* centromeric histone CID or the telomere capping protein HOAP, we conducted immunofluorescence (IF) on acid-free metaphase chromosome squashes. The B chromosomes are able to incorporate CID (Figure 3), confirming previous results (Bauerly *et al.* 2014). We observed HOAP localization at the ends of all the chromosomes, including the B chromosomes, leading us to conclude they are linear chromosomes with functional telomeres (Figure 3).

**Figure 3.**
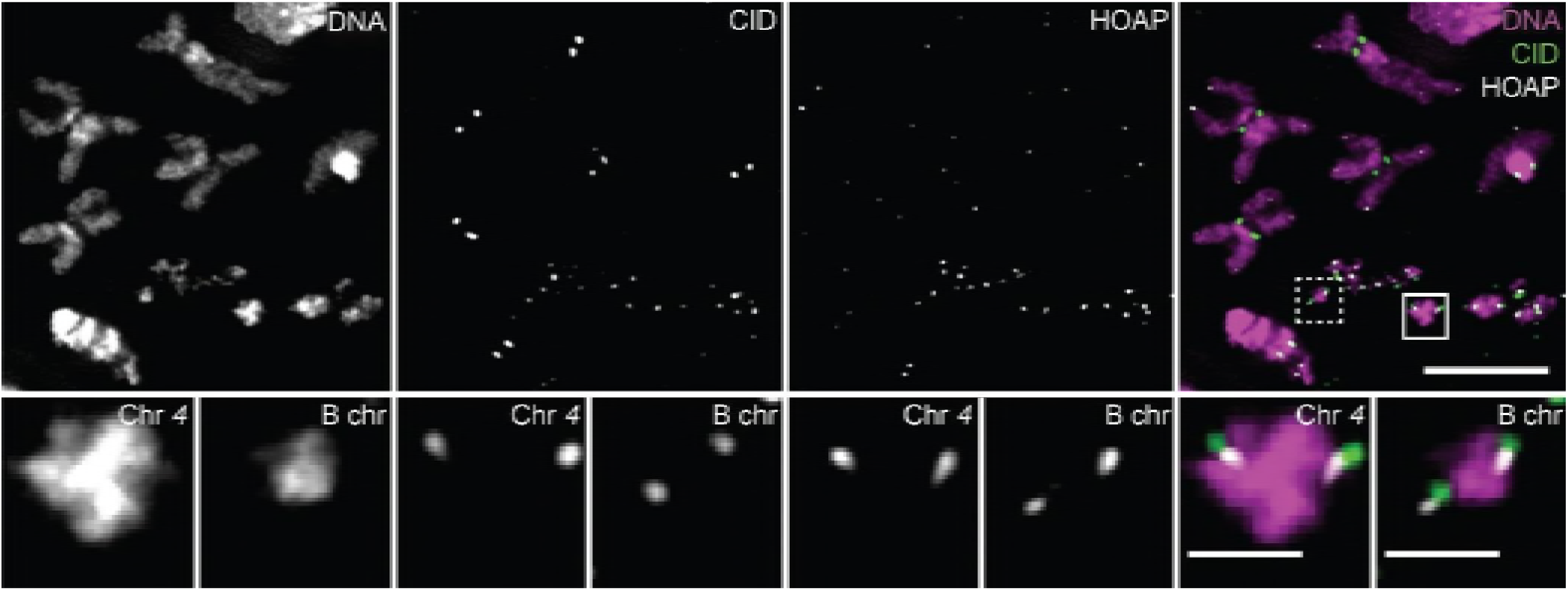
The B chromosomes have functional telomeres. Immunofluorescence with antibodies to CID (present at centromeres) and HOAP (present at telomeres) on metaphase chromosome spreads with B chromosomes. Magnified images are of Chromosome *4* (solid-line box in whole spread) and a B chromosome (dashed-line box in whole spread). Scale bar in whole spreads = 5 μm; scale bar in magnifications = 1 μm.

### Location and abundance of the *AAGAT* satellite repeat indicates the B chromosome is structurally an isochromosome

During our FISH analysis, we noticed that the number of fluorescent foci observed on Chromosome *4* with the *AAGAT* probe was never more than two (which we interpret as one focus per chromatid), whereas within the same metaphase, the B chromosomes could have as many as four foci. This observation is not dependent on degree of chromosome condensation as we do not treat the tissue with a cell cycle inhibitor prior to performing FISH (Figure 4A-C). If the additional FISH foci are due to the *AAGAT* satellite repeat block being split on the B chromosomes, then the intensity of the FISH signal should be similar to that of Chromosome *4*. However, if there is more of the *AAGAT* satellite repeat on the B chromosomes, then their FISH signal would be greater than that of Chromosome *4*. We therefore measured the intensity of the *AAGAT* FISH signal on individual B chromosomes and compared it to the signal on individual Chromosome *4s* within the same metaphase (see Materials and Methods). After analyzing 29 metaphases, we found a twofold difference in the *AAGAT* FISH probe signal intensity when comparing the B chromosomes to Chromosome *4* (Figure 4D).

**Figure 4.**
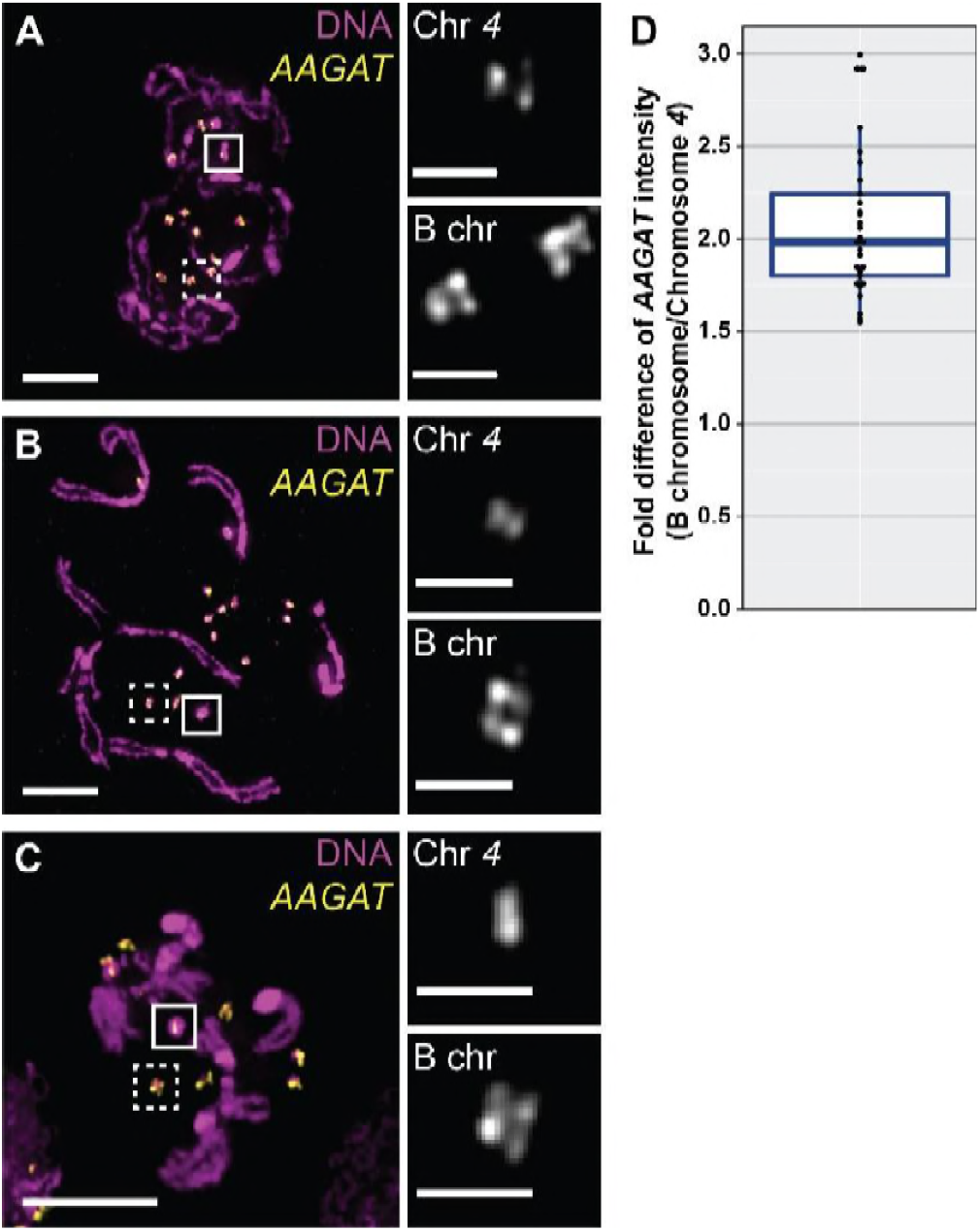
The B chromosomes have twice as much *AAGAT* sequence as Chromosome *4* as assayed by FISH. Chromosome spreads showing under-condensed (A), moderately condensed (B), and well-condensed (C) chromosomes. Magnified images of the *AAGAT* FISH signal on Chromosome *4* (in solid-line boxes) and the signal on the B chromosomes (in dashed-line boxes) illustrate how B chromosomes can have up to four *AAGAT* foci whereas Chromosome *4* has up to two. Scale bar in whole spreads = 5 μm; scale bar in magnifications = 1 μm. (D) Quantitative comparison of *AAGAT* FISH probe fluorescence signal indicates a twofold intensity difference of the B chromosome when compared to Chromosome *4*.

This intensity is twice what we would expect if the B chromosome was derived from a single chromosome fragment, leading us to reassess how the *D. melanogaster* B chromosome may have arisen from Chromosome *4*. We considered the reported structures of B chromosome in other species, such as the B chromosome in the characid fish *Astyanax scabripinnis*. This B chromosome appears to be an isochromosome that is thought to have formed after a centromeric misdivision event (Mestriner *et al.* 2000). Such an event separates the left arms from the right arms of a chromatid pair due to a break at the centromere; if the two identical arms fuse together, they simultaneously reconstitute the centromere and heal the broken chromosomes ends, resulting in an isochromosome. Focusing on the apparent duplication of the *AAGAT* satellite repeat on the B chromosomes as compared to Chromosome *4*, we propose the *D. melanogaster* B chromosome is an isochromosome that formed after a centromeric misdivision event on Chromosome *4* (see Discussion).

### Identification of a second B chromosome variant

Recently, we began examining the progenitor of the *mtrm^126^* stock that is retained in our laboratory. This stock carries a different null allele of *mtrm* caused by a P-element insertion upstream of the open reading frame, herein designated *mtrm^KG08051^* (Harris *et al.* 2003; Xiang *et al.* 2007). In a few of our samples from this stock, the metaphase chromosome spreads included a small DAPI-staining fragment that was reminiscent of a B chromosome. To determine if this DNA fragment had a similar repeat composition as the B chromosome, we performed FISH on metaphase spreads using probes for repeat sequences that are enriched on the B chromosome, revealing that this small DNA fragment carries each of them (Figure 5A, Figure S3). Since this DNA fragment was not present in every sample we examined, we speculated it may lack essential chromosome components such as a centromere and/or telomeres, thus affecting its ability to be efficiently transmitted to progeny. Therefore, we performed IF on metaphase spreads from the *mtrm^KG08051^* stock and found that this element carries both the CID and HOAP antigens, suggesting that it possesses both centromeres and telomeres, respectively (Figure 5B). Taken together, we believe this DNA fragment is the second B chromosome variant identified in *D. melanogaster* and will herein refer to it as the “B2” chromosome to distinguish it from the original B chromosome (now referred to as “B1”).

**Figure 5.**
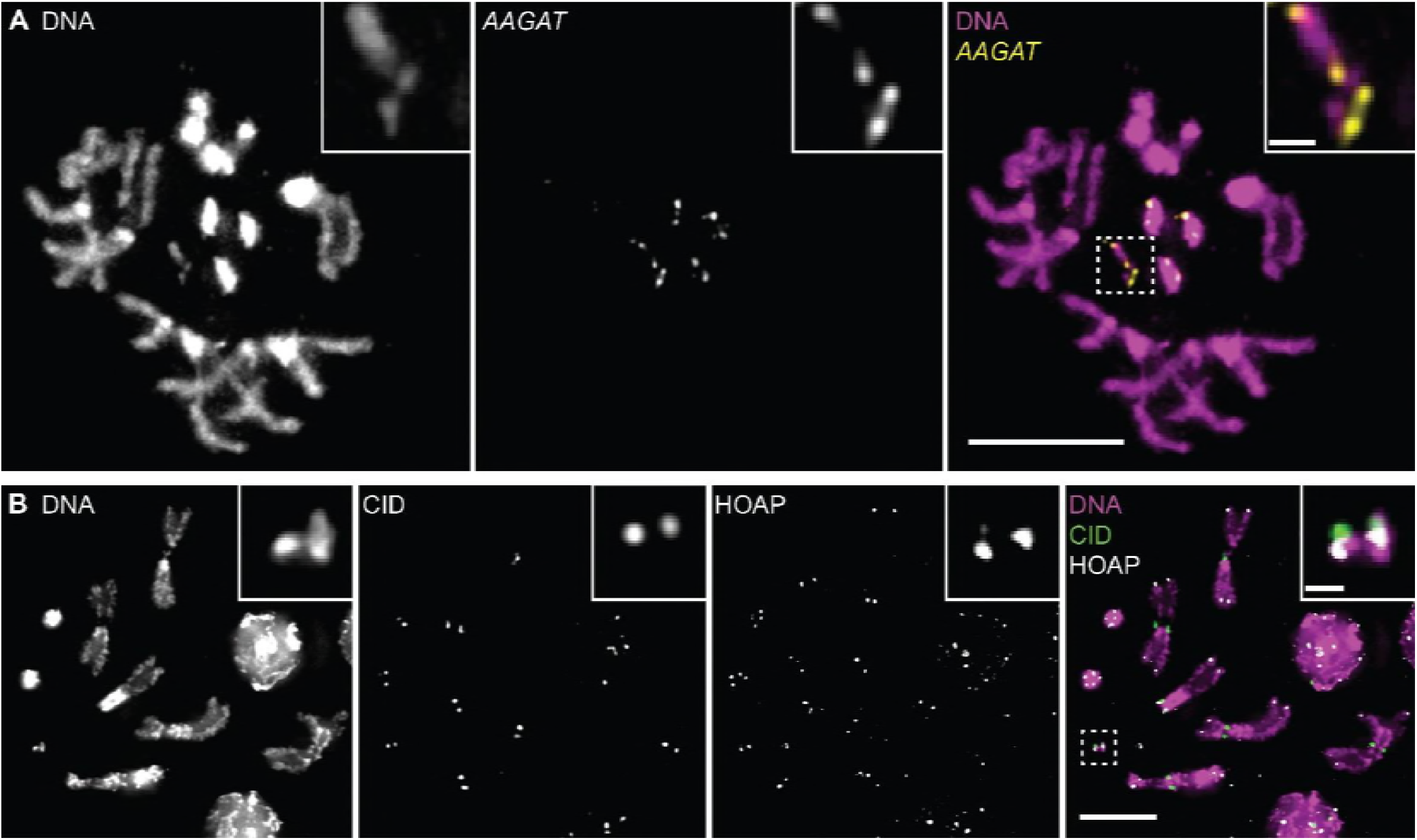
The B2 chromosome carries the *AAGAT* repeat and has centromeres and telomeres. (A) A FISH probe recognizing the *AAGAT* satellite repeat hybridizes to the B2 chromosome in metaphase chromosome spreads. Inset shows the magnification of two B2 chromosomes contained within the dashed-line box. Note: this larva carried three copies of Chromosome *4*, which is not uncommon in a *mtrm* null heterozygote (see text). (B) Immunofluorescence with antibodies to CID (present at centromeres) and HOAP (present at telomeres) on chromosome spreads in the *mtrm^KG08051^* stock that carries the B2 chromosome. Inset shows the magnification of a B2 chromosome contained within the dashed-line box. Scale bar in whole spreads = 5 μm; scale bar in magnifications (insets) = 0.5 μm.

Though the B1 and B2 chromosomes have a similar composition, there are some apparent cytological differences that exist between them. FISH experiments with the *AAGAT* probe showed that the B2 chromosome has fewer FISH foci—as well as less overall DAPI intensity—than does the B1 chromosome. To directly compare the two B chromosomes to each other as well as to the same Chromosome *4*, a male carrying the B1 chromosome was crossed to a female carrying the B2 chromosome. Progeny from this cross would possess both B chromosomes, enabling their direct comparison since they are within the same metaphase (Figure 6A). (We should note that the quantitative comparisons between the B1 chromosome and Chromosome *4* in Figure 4D were performed on the same metaphases, therefore the Chromosome *4* used for comparison here is identical.) Comparing the *AAGAT* FISH signal intensity on the B2 chromosome to Chromosome *4* demonstrates that they have an equal amount of FISH signal (Figure 6B, see Materials and Methods). This result is consistent with our observation that the B2 chromosome has fewer FISH foci than the B1 chromosome (Figure 6C). Since both B chromosome variants carry a similar suite of satellite repeats and are likely similar in their DNA composition, we used their DAPI staining intensity to relatively compare their DNA content. We found that the B2 chromosome is 65% as bright as the B1 chromosome, suggesting the B2 chromosome has less DNA content overall and is likely smaller in total length than the B1 chromosome (Figure 6D, see Materials and Methods). Due to these differences, we conclude that the B1 and B2 chromosomes are distinct and represent two variants of B chromosomes in *D. melanogaster*.

**Figure 6.**
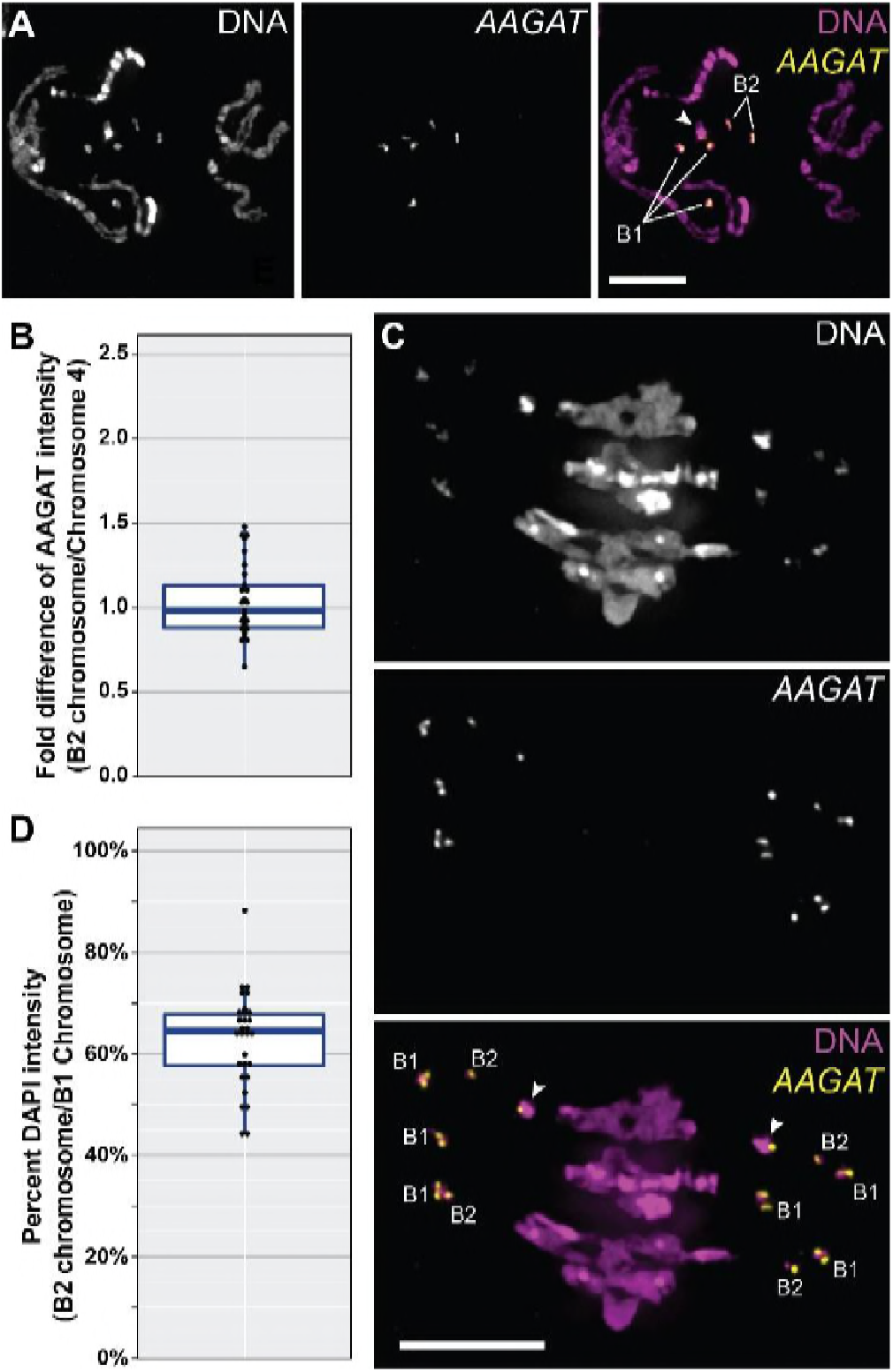
The B1 and B2 chromosome variants are distinct. (A) Metaphase chromosome spread with three copies of the B1 chromosome, two copies of the B2 chromosome, and a single copy of Chromosome *4* (denoted by an arrowhead) Note: this larva carried one copy of Chromosome *4*, which is not uncommon in a *mtrm* null heterozygote (see text). Scale bar = 5 μm (B) Quantitative comparison of *AAGAT* FISH probe fluorescence signal indicates the B2 chromosome has the same amount of signal as Chromosome *4*. (C) A cell in early anaphase was observed in the same tissue that produced (A). Here, as in metaphase chromosome spreads, the B1 chromosome is observed to have twice the *AAGAT* signal as the B2 chromosome. Scale bar = 5 μm (D) Quantitative comparison of DAPI staining intensity between the B1 and B2 chromosome. indicating the B2 chromosome has less DNA content than the B1 chromosome.

### Other stocks carrying the *mtrm^KG08051^* allele do not have B chromosomes and lack the *AAGAT* repeat

The *mtrm^KG08051^* stock carrying the B2 chromosome was originally a gift from the Bellen lab and has been in our laboratory collection since 2003 (Harris *et al.* 2003). Copies of this stock carrying the same *mtrm^KG08051^* null allele also exist in our institute stock collection as well as the Bloomington Drosophila Stock Center (herein referred to as *mtrm^SIMR1^*, *mtrm^SIMR2^*, and *mtrm^BDSC^*, respectively; see Table S1). To determine if any of these three lines also carry a B chromosome and at what frequency, we examined them cytologically and looked for small DAPI-staining fragments. Although previous chromosome spreads described here were made from larval neuroblast tissue, for this analysis we chose to use ovarian tissue from adult females in order to quickly assess how many, if any, breeding adults carried B chromosomes within each of the stocks. We began by examining the *mtrm^KG08051^* line that we know carries the B2 chromosome in order to determine its frequency within the stock. We found that nine out of 25 adult females carried one copy of the B2 chromosome and two females carried two copies of the B2 chromosome, indicating its frequency within the stock is very low (Figure S4A). We then examined females from the other three stocks using the same technique and included the *AATAT* FISH probe in our chromosome spread processing since it is enriched on both the B1 and B2 chromosomes. Interestingly, after examining at least 30 females from each stock, we did not detect a supernumerary DAPI-staining fragment (Figure S4B-E). We therefore conclude that the *mtrm^SIMR1^*, *mtrm^SIMR2^*, and *mtrm^BDSC^* lines do not carry B chromosomes.

During the initial examination of the *mtrm^SIMR1^*, *mtrm^SIMR2^*, and *mtrm^BDSC^* lines, we used the *AAGAT* FISH probe in our chromosome spread processing. We ultimately switched to the *AATAT* probe for our analyses, however, since we were unable to detect *AAGAT* FISH signal in the other three lines. This result was puzzling for two reasons. First, though these three stocks were acquired from separate collections, they are genotypic copies of our laboratory *mtrm^KG08051^* stock that clearly carries the *AAGAT* satellite repeat (Figure 7A) and therefore we expected them to be similar. Second, since the *AAGAT* satellite repeat was abundant in two separate genomic studies (Wei *et al.* 2014; Talbert *et al.* 2018), we were surprised to not detect any blocks of *AAGAT* satellite repeat in these three stocks. To validate our original observation, we repeated the FISH experiment on neuroblast chromosome spreads from males (in order to also assay the *Y* chromosome) using both the *AAGAT* and *AATAT* probes simultaneously. Consistent with our initial results, we observed FISH signal with the *AATAT* probe but not the *AAGAT* probe in these three stocks (Figure 7B-D). This result confirms that the lack of *AAGAT* signal we observe is due to polymorphism in the amount of *AAGAT* satellite repeat carried by the variants of Chromosome *4* in these stocks.

**Figure 7.**
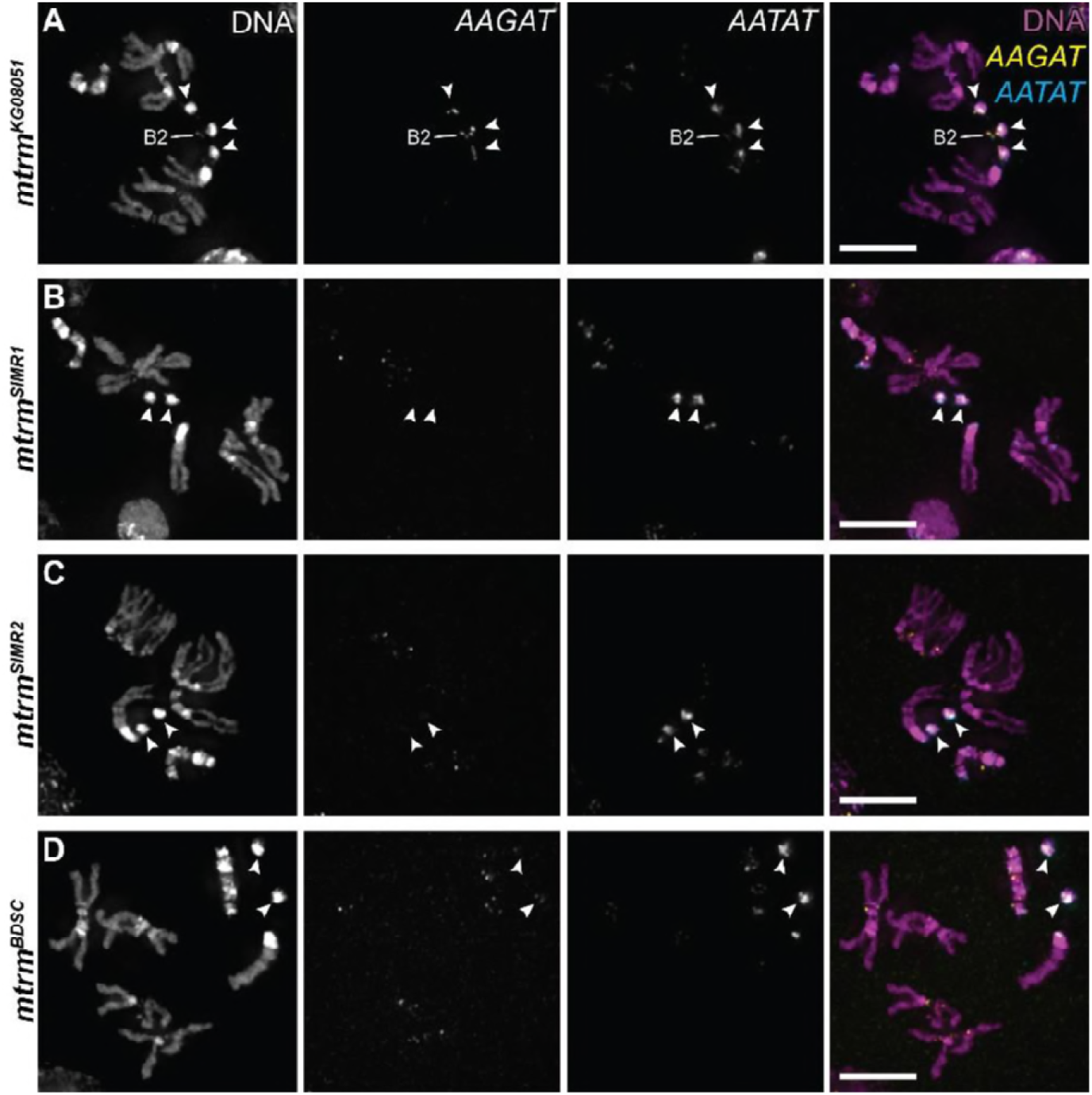
Other *mtrm^KG08051^* stocks do not carry B chromosomes and are polymorphic for the *AAGAT* satellite repeat. (A) The laboratory version of the *mtrm^KG08051^* stock carries the B2 chromosome and has the *AAGAT* repeat on Chromosome *4* (denoted by arrowheads) as assessed by FISH. Note: this larva carried three copies of Chromosome *4*, which is not uncommon in a *mtrm* null heterozygote (see text). The other *mtrm^KG08051^* lines *mtrm^SIMR1^* (B), *mtrm^SIMR2^* (C), and *mtrm^BDSC^* (D) do not possess B chromosomes, nor do they carry the *AAGAT* satellite repeat (Chromosome *4* denoted by arrowheads). Brightness and contrast were adjusted for the DNA (DAPI) channel only (see Materials and Methods). Scale bar = 5 μm.

## DISCUSSION

In the century since the first description of a supernumerary chromosome, the existence of B chromosomes has been well documented in several species spanning a diverse range of taxa. Despite their widespread prevalence, only a handful of systems have possessed tools to enable an in-depth molecular and genomic analysis of their B chromosome(s) (López-León *et al.* 1994; Han *et al.* 2007; Banaei-Moghaddam *et al.* 2012; Kour *et al.* 2014; Ellis *et al.* 2015; Huang *et al.* 2016; Aldrich *et al.* 2017; Ruiz-Ruano *et al.* 2018). The appearance of B chromosomes in an established model animal system such as *D. melanogaster* enables us to use established molecular tools to study the anatomy of a newly formed chromosome and quickly ascertain its origin. By isolating and deep-sequencing the B1 chromosome, we were able to determine that it originated from the *D. melanogaster* genome, lacks known euchromatic material, and is comprised of several heterochromatic elements.

One of the heterochromatic elements we identified is the *AAGAT* satellite repeat, which has been recently reported but was not characterized (Wei *et al.* 2014; Talbert *et al.* 2018). The *AAGAT* repeat was found to be the fourth-most abundant 5mer repetitive sequence in a whole-genome analysis of several Drosophila lines (Wei *et al.* 2014), and is likely positioned near a centromere as indicated by its enrichment in CENP-A ChIP experiments (Talbert *et al.* 2018). Though Talbert *et al.* (2018) proposed it as a candidate for the centromeric satellite repeat on the *Y* chromosome, we did not detect *AAGAT* sequence on the *Y* chromosome via FISH in any of our stocks, including the stocks that lacked the *AAGAT* satellite repeat on Chromosome *4* (Figure 7). Rather, we have shown here that this repeat appears to be uniquely localized to Chromosome *4* and to both B chromosomes. Together, its evident abundance at centromeres (Talbert *et al.* 2018) and our chromosome cytology of Chromosome *4* and the B chromosomes indicate the *AAGAT* satellite sequence is likely associated with the centromere on Chromosome *4*.

Based on the *AAGAT* satellite repeat cytology, the structure of the B1 chromosome is most consistent with that of an isochromosome formed from Chromosome *4* (Figure 8). It is likely comprised of the shorter, left arm of Chromosome *4* since the right arm carries euchromatic material, which has been shown both previously (Bauerly *et al.* 2014) and in this study to be absent from the B chromosomes. One well-documented mechanism of isochromosome formation is through a centromeric misdivision event (Rhoades 1933; Upcott 1937; Darlington 1939; Carlson 1970; Kaszás and Birchler 1996). The resulting chromosome fragments can fuse to become an isochromosome, thus eliminating the broken ends and potentially reconstituting a full centromere. It has been demonstrated in wheat that the formation of isochromosomes from univalent (unpaired) chromosomes through a centromeric misdivision event is quite frequent during meiotic divisions (Sears 1952; Steinitz-Sears 1966; Friebe *et al.* 2005; Lukaszewski 2010). Such an event may have the opportunity to occur more often in a *mtrm* mutant background due to an increased frequency of Chromosome *4* aneuploidy within the stock, a result of its frequent missegregation during female meiosis (Harris *et al.* 2003; Xiang *et al.* 2007; Bonner *et al.* 2013). Females carrying an extra copy of Chromosome *4* would produce a univalent during meiosis, which may be more susceptible to centromeric misdivision. Additionally, B chromosomes carried by other species have been shown to be isochromosomes that arose from one of the essential A chromosomes (John and Hewitt 1965; Vicente *et al.* 1996; Mestriner *et al.* 2000; Dhar *et al.* 2002; Poletto *et al.* 2010). Indeed, as suggested by Battaglia (1964), misdivision and isochromosome formation is likely a recurring mechanism for B chromosome creation.

**Figure 8.**
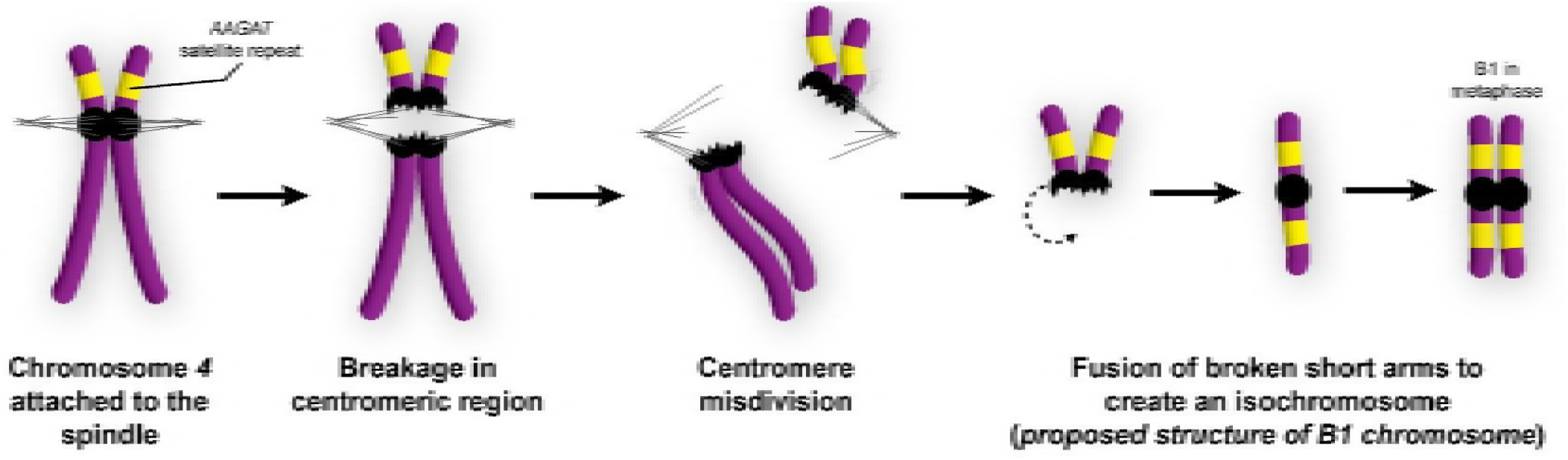
Model for B1 chromosome formation in *D. melanogaster*. FISH analysis of the *AAGAT* repetitive sequence (depicted in yellow) indicates the B1 chromosome is an isochromosome (see Figure 4 and text). Such a chromosome may form after the misdivision of the centromere on Chromosome *4*, leaving the fragments from the short, left arm to fuse and become the B1 chromosome.

We also identified a second, smaller B chromosome variant (B2) in a separate laboratory stock carrying a different null allele of *mtrm*. Though the B2 chromosome carries the same suite of repeats as the B1 chromosome, has both telomeric and centromeric components, and was found in a similar mutant background, the copy number and frequency of the B2 chromosome in its endogenous stock (*mtrm^KG08051^*) is substantially lower than is characteristic of the B1 chromosome copy number in its endogenous stock (*mtrm^126^*). This observation may be attributed to the variation in genetic backgrounds (see Table S1) or the difference in the size of the two B chromosomes. Periodic testing of this stock will enable us to use established methods to determine if the prevalence of this new B chromosome is fluctuating and, if so, at what rate.

Though the B1 chromosome structure is consistent with an isochromosome, we are unsure what the structure of the B2 chromosome may be. Since the DAPI intensity of the B2 chromosome is 65% that of the B1 chromosome (Figure 6D), we favor a model where the B2 chromosome arose after a centromeric misdivision event that encompassed a larger portion of the centromeric region, followed by the acquisition of a telomere instead of the joining of the broken fragments. We are aware, however, that due to the shared histories of the stocks each B chromosome was found in, we cannot establish whether each B chromosome arose independently, or if one was formed from another. Though its size and prevalence preclude us from being able to extract it from a pulsed-field gel experiment, newer sequencing technologies that produce extremely long reads may enable us to circumvent this limitation. Variations such as single-nucleotide polymorphisms, small insertions and deletions, or transposable element positions that differ between the B chromosomes and Chromosome *4* may be used to distinguish the order of origin of the B chromosomes and help determine what, if any, constraints exist on B chromosome evolution. Additionally, long-read sequencing of the satellite repeat regions will produce a more accurate analysis of their composition, allowing us to track their changes over time and in various genetic backgrounds.

Our results strongly indicate the B chromosomes originated from Chromosome *4*, however we presently are unaware as to how often they arise and how long they can be maintained. To this end, we are curious what may make the original *mtrm^126^* stock (with B1 chromosomes) and the single *mtrm^KG08051^* stock (with B2 chromosomes) unique. Though all five stocks we examined carry a null allele of *mtrm* (Table S1), only the stocks with B chromosomes also have the *AAGAT* satellite repeat on Chromosome *4* (as assessed by FISH on metaphase chromosome spreads, Figure 7). We are intrigued as to what role, if any, this repetitive sequence plays in the formation and/or maintenance of a B chromosome.

Ultimately, a deeper understanding of the structure and formation of the *D. melanogaster* B chromosomes may provide insight into the etiology of similar chromosomes that can be detrimental to human health. Small supernumerary marker chromosomes (sSMCs) in humans have many properties that are similar to B chromosomes (Fuster *et al.* 2004; Liehr *et al.* 2008). These sSMCs arise from the A chromosomes and are present in 1 out of ~2,300 newborns (Liehr and Weise 2007). Many are connected to specific syndromes, such as Emanuel, cat eye, and Pallister-Killian syndromes, or are associated with intellectual disabilities and infertility (Liehr 2012; Armanet *et al.* 2015; Jafari-Ghahfarokhi *et al.* 2015; Wang *et al.* 2015). Improved molecular characterization is beginning to reveal more about the genomic organization of supernumerary chromosomes (Breman *et al.* 2011; Sun *et al.* 2017). A subset of sSMCs are isochromosomes derived from two of the short arms of a single chromosome after an apparent centromeric misdivision event (de la Chapelle 1982; Bugge *et al.* 2004; Jafari-Ghahfarokhi *et al.* 2015). Isochromosomes are also observed in various cancers, most notably testicular germ cell tumors where isochromosome 12p is a well-established hallmark and is believed to be a triggering event for invasive growth (Litchfield *et al.* 2016; Mitelman *et al.* 2018). Continued investigation of the mechanisms behind supernumerary chromosome formation, retention, and transmission using *D. melanogaster* B chromosomes as a model system will thus be of interest to a wide range of research areas.

## ACKNOWLEDGEMENTS

We would like to thank the Hawley Laboratory for their support and helpful comments on the manuscript. We also thank Rhonda Egidy, Anoja Perera, and the Molecular Biology core at SIMR for expert help with DNA sequencing. The rabbit anti-HOAP serum was a generous gift from Yikang S. Rong at Sun Yat-sen University in China, and we thank him for sharing it with us. RSH is supported by the Stowers Institute for Medical Research and is an American Cancer Society Research Professor.

